# Metabolic effects of aripiprazole and olanzapine multiple-dose treatment in healthy volunteers. Association with pharmacogenetics

**DOI:** 10.1101/2020.07.29.226209

**Authors:** Dora Koller, Susana Almenara, Gina Mejía, Miriam Saiz-Rodríguez, Pablo Zubiaur, Manuel Román, Dolores Ochoa, Marcos Navares-Gómez, Elena Santos-Molina, Elena Pintos-Sánchez, Francisco Abad-Santos

**Affiliations:** Clinical Pharmacology Department, Hospital Universitario de La Princesa, Instituto Teófilo Hernando, School of Medicine, Universidad Autónoma de Madrid, Instituto de Investigación Sanitaria La Princesa (IP), Madrid, Spain; UICEC Hospital Universitario de La Princesa, Platform SCReN (Spanish Clinical Research Network), Instituto de Investigación Sanitaria La Princesa (IP), Madrid, Spain; Research Unit, Fundación Burgos por la Investigación de la Salud, Hospital Universitario de Burgos, Burgos, Spain

**Author notes:** Corresponding author Dr. Francisco Abad-Santos, Clinical Pharmacology Department, Hospital Universitario de La Princesa, Diego de León 62., 28006 Madrid, Spain.

**Keywords:** aripiprazole, olanzapine, metabolism, pharmacogenetics, pharmacokinetics

## Abstract

**Background:** Aripiprazole and olanzapine are atypical antipsychotics. Both drugs can induce metabolic changes, however, the metabolic side effects produced by aripiprazole are more benign.

**Objectives:** To evaluate if aripiprazole and olanzapine alter prolactin levels, lipid and glucose metabolism and hepatic, hematological, thyroid and renal function.

**Methods:** Twenty-four healthy volunteers received 5 daily oral doses of 10 mg aripiprazole and 5 mg olanzapine in a crossover randomized clinical trial and were genotyped for 51 polymorphisms in 17 genes by qPCR. Drug plasma concentrations were measured by LC-MS. The biochemical and hematological analyses were performed by enzymatic methods.

**Results:** Olanzapine induced hyperprolactinemia but not aripiprazole. *DRD3* Ser/Gly and *ABCB1* rs10280101, rs12720067 and rs11983225 polymorphisms and CYP3A phenotype had an impact on plasma prolactin levels. C-peptide concentrations were higher after aripiprazole administration and were influenced by *COMT* rs4680 and rs13306278 polymorphisms. Olanzapine and the *UGT1A1* rs887829 polymorphism were associated with elevated glucose levels. CYP3A poor metabolizers had increased insulin levels. Triglyceride concentrations were decreased due to olanzapine and aripiprazole treatment and were variable based on CYP3A phenotypes and the *APOC3* rs4520 genotype. Cholesterol levels were also decreased and depended on *HTR2A* rs6314 polymorphism. All hepatic enzymes, platelet and albumin levels and prothrombin time were altered during both treatments. Additionally, olanzapine reduced the leucocyte count, aripiprazole increased free T4 and both decreased uric acid concentrations.

**Conclusions:** Short term treatment with aripiprazole and olanzapine had a significant influence on the metabolic parameters. However, it seems that aripiprazole provokes less severe metabolic changes.

## 1. Introduction

Antipsychotic drugs are indicated for the treatment of schizophrenia and other psychotic disorders (schizoaffective disorder, delusional disorder and bipolar affective disorder, among others). Habitually, they are categorized as first-generation (typical) antipsychotics (FGAs) or second-generation (atypical) antipsychotics (SGAs) ^1^. Olanzapine (OLA) acts as an antagonist at dopamine D2 receptors (DRD2) and at serotonin (5HT) 1A, 2A and 2C receptors similarly to other second-generation antipsychotics ^2^. Contrastively, aripiprazole (ARI) acts as a partial agonist at DRD2 and dopamine D3 receptors (DRD3) and at 5HT1A and 5HT2C receptors and has antagonistic activity at 5HT2A receptors. Therefore, ARI possesses a unique mechanism of action compared to other SGAs ^3^.

ARI is extensively metabolized by cytochrome P450 (CYP) isoforms CYP3A4 and CYP2D6. The main active metabolite – dehydro-aripiprazole (DARI) – represents 40% of the parent drug in steady state ^4^. OLA is metabolized predominantly by direct glucuronidation via the UDP-glucuronosyltransferase (UGT) enzyme family and CYP1A2 and to a lesser extent by CYP2D6 and CYP3A4 ^5^.

Several atypical antipsychotics – including OLA – cause plasma prolactin level elevation ^6^. Normally, while switching the therapy from OLA to ARI mean prolactin levels decrease significantly even after one week of treatment and are maintained forth ^7^. Nevertheless, ARI can also cause mild prolactin elevation in less than 5% of patients ^6^. OLA is a D2 receptor antagonist and induce hyperprolactinaemia via inhibition of dopamine action at D2 receptors in the hypothalamus, where prolactin secretion is regulated ^6^. On the contrary, serotonin stimulates prolactin secretion probably via stimulation of prolactin-releasing factors ^8^. ARI acts as a functional antagonist under hyperdopaminergic conditions while it acts as a functional agonist under hypodopaminergic conditions at dopamine D2 receptors. D2 receptor stimulation provokes a suppression on prolactin secretion, therefore ARI’s high D2 receptor occupancy does not induce hyperprolactinemia in the majority of subjects ^9^.

*DRD2* polymorphisms affected prolactin secretion induced by OLA in healthy volunteers after a single dose administration and in schizophrenic patients under chronic treatment ^10,11^. Nevertheless, prolactin concentrations were not affected by *DRD2* polymorphisms and CYP2D6 phenotype after ARI administration in schizophrenic patients ^12^. On the contrary, an association was found between prolactin levels and CYP2D6 phenotype and *HTR2C* polymorphisms in healthy volunteers ^13^.

Previous clinical trials with schizophrenic patients demonstrated that ARI has more benign side effect profile – regarding weight gain, blood sugar level and lipid profile – as compared to OLA in short-term treatment. Weight gain was observed more frequently in OLA-treated patients when compared with ARI. Mean serum triglyceride, blood glucose and cholesterol levels in patients treated with OLA were higher than in patients treated with ARI ^14,15^. Additionally, OLA was associated with significantly increased glucose levels compared to placebo and with a significantly greater change in the glucose levels compared to other antipsychotics ^16^. In addition, when comparing lean mice and others on high-fat diet, OLA induced hyperglycemia and therefore systemic insulin resistance ^17^.

The aim of the current study was to evaluate the effect of ARI and OLA after multiple dose administration to healthy volunteers on lipid and glucose metabolism, hepatic, haematological and thyroid performance and prolactin levels and its correlation with various factors including sex, plasma drug concentrations and selected genetic polymorphisms.

## 2. Results

### 2.1. Demographic characteristics

The study was performed between June 2018-April 2019 including recruitment and follow-up visits. Demographic data are shown in *Table* S1. Ten subjects were Caucasian and 14 were Latin. Average age was similar between males and females (p = 0.204). Males had greater weight and height than females (p < 0.001), however, the BMI values did not differ significantly (p = 0.798).

### 2.2. Genotype frequencies

The genotype frequencies of the analysed genes are shown in *Table* S2. *HTR2C* rs3813929 and rs518147, *ABCB1* rs4728709 and *COMT* rs13306278 were not in Hardy-Weinberg equilibrium (p ≤ 0.05). The rest of the polymorphisms were in Hardy-Weinberg equilibrium (p ≥ 0.05).

Genotype frequencies of *ABCB1* C1236T, G2677T/A, 10276036 and rs4148737 and *HTR2C* rs518147 polymorphisms were significantly different between males and females (*Table* S2).

### 2.3. Pharmacokinetic analysis

Mean and standard deviation (SD) of ARI, DARI and OLA pharmacokinetic parameters are shown in *Table* 1. Females had lower DARI/ARI ratio than males (p = 0.046). The remaining pharmacokinetic parameters were not statistically different between the two sexes. Associations between pharmacokinetic parameters and polymorphisms were reported previously ^18^.

### 2.4. Prolactin concentrations and their relationship with pharmacogenetics

OLA caused a significant elevation in prolactin levels (p < 0.001, partial eta squared (η_p_^2^) = 0.474) (*Figure* 1). Males had lower prolactin levels than females, however, the extent of the increment did not differ between them. Additionally, a significant interaction was found between OLA C_max_ and prolactin levels (p = 0.006, η_p_^2^ = 0.168). Moreover, volunteers carrying the *DRD3* Gly carriers had significantly higher prolactin concentrations than volunteers with the Ser/Ser genotype (p = 0.036, η_p_^2^ = 0.121).

Compared to OLA, ARI did not elevate prolactin levels. On the contrary, a tendency of decrease was observed but it did not reach the significant level (p = 0.052) (*Figure* 1). Additionally, CYP3A PMs had higher prolactin concentrations during ARI treatment compared to IMs and EMs (p = 0.001, η_p_^2^ = 0.226). *ABCB1* rs10280101 A/A, rs12720067 C/C and rs11983225 T/T subjects had significantly higher prolactin concentrations compared to C, T and C allele carriers (p = 0.037, η_p_^2^ = 0.123). However, when analysing *ABCB1* haplotypes, this association could not be detected.

**Figure 1.**
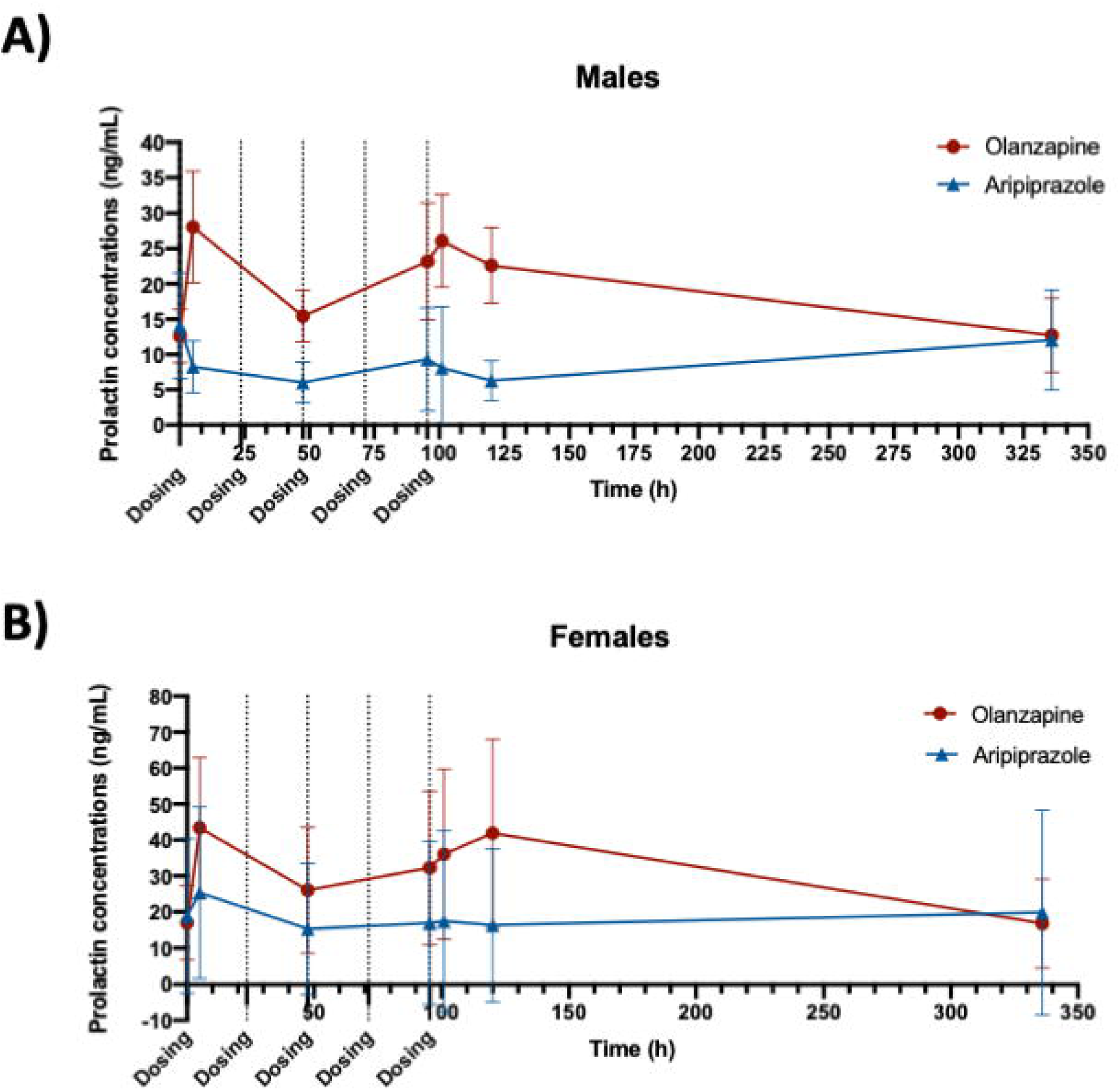
Prolactin concentrations during the administration of 5 daily doses of aripiprazole 10 mg and olanzapine 5 mg tablets. The results are shown in mean ± SD. ***Figure* 1A**. Prolactin concentrations versus time in males treated with aripiprazole and olanzapine. ***Figure* 1B**. Prolactin concentrations versus time in females treated with aripiprazole and olanzapine.

The same genetic associations were found in males and females. Prolactin levels were significantly higher during OLA treatment compared to ARI (p < 0.0001, η_p_^2^ = 0.356). Prolactin levels were outside of the recommended range (males: 2-18 ng/mL; females: 3-30 ng/mL) during OLA treatment in 9 (75%) males and 9 (75%) females ^19^.

### 2.5. Glucose metabolism and its relationship with pharmacogenetics

C-peptide concentrations were significantly higher after completing ARI treatment compared to its initial levels (p = 0.030, η_p_^2^ = 0.205) (*Table* 2). Additionally, the increase of C-peptide levels was greater in *COMT* rs4680 G/G subjects and rs13306278 T carriers compared to A carriers and C/C homozygotes, respectively (p = 0.010, η_p_^2^ = 0.289; p < 0.001, η_p_^2^ = 0.535, respectively). This association could not be detected when analysing the COMT phenotype. Moreover, although the insulin levels only tended to increase after ARI administration (p = 0.073), *BDNF* rs6265 C/C subjects had greater increment compared to the other genotypes (p = 0.040, η_p_^2^ = 0.237). Likewise, DARI AUC_last_ was indirectly proportional with the changes in insulin levels (p = 0.045, η_p_^2^ = 0.228).

After completing OLA treatment, the 1 h and 2 h glucose levels after performing GTT were higher compared to the measurements on the first day (p = 0.007, ηp2 = 0.213) (*Table* 2). In addition, these changes were dependent on the *UGT1A1* rs887829 genotype: C/C homozygote subjects had significantly higher glucose concentrations in GTT both after 1 h and 2 h than the T allele carriers (p = 0.014, η_p_^2^ = 0.186). Moreover, this polymorphism was additionally related to higher increase in glucose levels in C/C subjects compared to the T allele carriers (p = 0.013, η_p_^2^ = 0.258). Additionally, the insulin levels of CYP3A PMs incremented more compared to EMs and IMs (p = 0.029, η_p_^2^ = 0.217). Moreover, OLA administration increased the C-peptide/insulin ratio (p = 0.044, η_p_^2^ = 0.196).

**Table 1.** Pharmacokinetic parameters of aripiprazole, dehydro-aripiprazole and olanzapine after administration of 5 multiple doses.

| **ARIPIPRAZOLE** | **All (n = 24)** | **Males (n = 12)** | **Females (n = 12)** |
| --- | --- | --- | --- |
| **AUC<sub>INF</sub> (ng·h/mL)** | 11102.4 ± 8234.0 | 7789.1 ± 4071.5 | 14415.7 ± 10061.4 |
| **C<sub>MAX</sub> (ng/mL)** | 137.94 ± 45.92 | 129.6 ± 47.4 | 146.3 ± 44.9 |
| **DEHYDRO-ARIPIPRAZOLE** |  |  |  |
| **AUC<sub>24H</sub> (ng·h/mL)** | 5149.8 ± 1628.6 | 4721.3 ± 1670.3 | 5578.3 ± 1534.8 |
| **C<sub>MAX</sub> (ng/mL)** | 34.88 ± 8.45 | 35.62 ± 9.62 | 34.13 ± 7.45 |
| **ARIPIPRAZOLE + DEHYDRO-ARIPIPRAZOLE** |  |  |  |
| **AUC<sub>24H</sub> (ng·h/mL)** | 14596.1 ± 6639.1 | 11883.5 ± 4788.9 | 17308.6 ± 7292.2 |
| **C<sub>MAX</sub> (ng/mL)** | 172.82 ± 48.74 | 165.27 ± 53.98 | 180.38 ± 43.93 |
| **DEHYDRO-ARIPIPRAZOLE/ ARIPIPRAZOLE** |  |  |  |
| **AUC RATIO** | 0.64 ± 0.25 | 0.74 ± 0.27 | 0.54 ± 0.20* |
| **C<sub>MAX</sub> RATIO** | 0.27 ± 0.08 | 0.29 ± 0.07 | 0.25 ± 0.09 |
| **OLANZAPINE** |  |  |  |
| **AUC<sub>INF</sub> (ng·h/mL)** | 1289.4 ± 370.1 | 1142.7 ± 291.2 | 1436.2 ± 393.1 |
| **C<sub>MAX</sub> (ng/mL)** | 19.10 ± 4.77 | 18.35 ± 4.04 | 19.85 ± 5.48 |

\*P ≤ 0.05 versus males.
Values are shown as mean ± SD.
Abbreviations: C<sub>max</sub>: maximum plasma concentration; AUC<sub>inf</sub>: area under the curve from zero to infinity; AUC<sub>24h</sub>: area under the curve over 24h; AUC ratio: dehydro-aripiprazole/aripiprazole AUC ratio; C<sub>max</sub> ratio: dehydro-aripiprazole/aripiprazole C<sub>max</sub> ratio.

**Table 2.** C-peptide, insulin, haemoglobin A1c and glucose levels during aripiprazole and olanzapine multiple dose treatment.

| ARIPIPRAZOLE |  |  |  |  |  |  |
| --- | --- | --- | --- | --- | --- | --- |
|  | Day 1 |  |  | Day 6 |  |  |
| C-peptide (ng/mL)* | 1.60 ± 0.35 |  |  | 3.50 ± 3.85 |  |  |
| Insulin (mcU/mL) | 7.84 ± 2.37 |  |  | 9.61 ± 4.37 |  |  |
| C-peptide/insulin ratio | 0.22 ± 0.06 |  |  | 0.24 ± 0.09 |  |  |
| HbA1c (%) | 5.30 ± 0.22 |  |  | 5.28 ± 0.21 |  |  |
|  | Screening |  | Day 3 | Day 6 |  | Day 15 |
| Glucose (mg/dL) | 80.33 ± 6.64 |  | 79.71 ± 6.49 | 80.87 ± 9.01 |  | 79.17 ± 5.84 |
|  | Day 1 |  |  | Day 6 |  |  |
|  | 0 h | 60 min | 120 min | 0 h | 60 min | 120 min |
| GTT (mg/dL) | 81.79 ± 7.79 | 103.38 ± 35.63 | 76.00 ± 13.40 | 84.54 ± 8.82 | 122.04 ± 31.80 | 97.38 ± 21.56 |
| OLANZAPINE |  |  |  |  |  |  |
|  | Day 1 |  |  | Day 6 |  |  |
| C-peptide (ng/mL) | 1.69 ± 0.51 |  |  | 2.60 ± 2.29 |  |  |
| Insulin (mcU/mL) | 8.19 ± 2.31 |  |  | 10.40 ± 12.76 |  |  |
| C-peptide/insulin ratio* | 0.21 ± 0.03 |  |  | 0.25 ± 0.08 |  |  |
| HbA1c (%) | 5.36 ± 0.23 |  |  | 5.07 ± 1.09 |  |  |
|  | Screening |  | Day 3 | Day 6 |  | Day 15 |
| Glucose (mg/dL) | 79.96 ± 7.52 |  | 79.38 ± 6.50 | 79.14 ± 8.05 |  | 80.71 ± 6.22 |
|  | Day 1 |  |  | Day 6 |  |  |
|  | 0 h | 60 min | 120 min | 0 h | 60 min | 120 min |
| GTT (mg/dL)* | 81.88 ± 6.73 | 104.08 ± 33.81 | 89.38 ± 20.98<sup>#</sup> | 79.71 ± 8.65 | 124.21 ± 38.07 | 101.96 ± 28.51 |

\*P ≤ 0.05. <sup>#</sup>p ≤ 0.05 compared to aripiprazole. Values are shown as mean ± SD. Abbreviations: HbA1c: haemoglobin A1c, GTT: glucose tolerance test.

On first day’s GTT test, glucose levels were higher in OLA-treated subjects compared to ARI (p = 0.011, ηp2 = 0.131), however, ARI showed the same tendency. However, the increment in glucose levels after GTT and the increment of C-peptide levels did not differ between ARI and OLA (see *Table* 2). No changes were detected in HbA1c.

No differences were found between males and females nor in C-peptide, insulin, glucose and GTT levels neither in the genetic associations. No levels were outside of the normal range.

### 2.6. Weight and lipid metabolism and their relationship with pharmacogenetics

During ARI treatment, volunteers’ weight decreased significantly (p < 0.0001, η_p_^2^ = 0.301). On the contrary, a tendency of weight gain was observed during OLA treatment, but it did not reach the significant level (p = 0.120) (*Figure* 2). Additionally, a significant difference was found when comparing weight changes between ARI and OLA treatment (p < 0.001, η_p_^2^ = 0.301). Moreover, *HTR2C* rs1414334 C/C subjects gained significantly more weight compared to T carriers (p = 0.002; η_p_^2^ = 0.196).

Triglyceride levels linearly decreased due to ARI and OLA administration (p = 0.009, η_p_ ^2^ = 0.177; p = 0.047, η_p_^2^ = 0.125, respectively) (*Figure* 2). No significant difference was found in the extent of this decrease between ARI and OLA (p = 0.593). Additionally, ARI C_max_, DARI C_max_ and ARI + DARI C_max_ were inversely proportional to triglyceride levels (p = 0.003, η_p_^2^ = 0.203; p < 0.001, η_p_ ^2^ = 0.327; p < 0.001, η_p_ ^2^ = 0.258, respectively). Moreover, CYP3A PMs had significantly greater decrease in triglyceride levels during ARI treatment compared to the other phenotypes (p < 0.001, η_p_^2^ = 0.296). Furthermore, *APOC3* rs4520 C/C homozygotes had lesser decrease in triglyceride concentrations after OLA administration than T allele carriers (p = 0.018, η_p_^2^ = 0.162).

**Figure 2.**
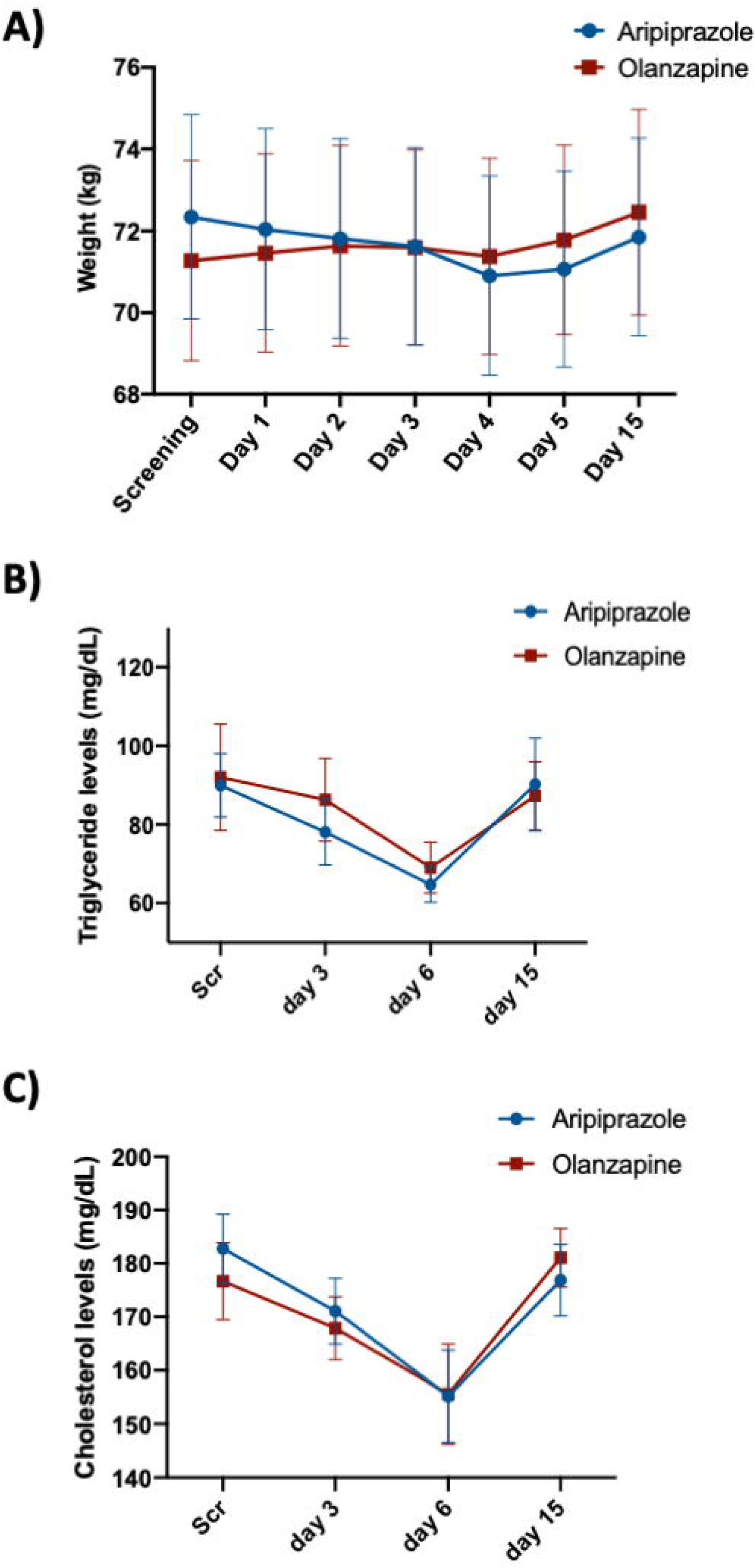
Weight and triglyceride and cholesterol concentrations during the administration of 5 daily doses of aripiprazole 10 mg and olanzapine 5 mg tablets. The values are shown as mean ± SD. ***Figure* 2A**. Changes in weight versus time. ***Figure* 2B**. Triglyceride concentrations versus time. ***Figure* 2C**. Cholesterol concentrations versus time.

Likewise, total cholesterol levels diminished significantly during ARI and OLA treatment (p = 0.002, η_p_ ^2^ = 0.250; p = 0.004, η_p_ ^2^ = 0.209, respectively) (*Figure* 2). No significant difference was found between ARI and OLA in the extent of this reduction (p = 0.241). Moreover, *HTR2A* rs6314 C/C subjects had higher cholesterol concentrations during ARI treatment compared to T allele carriers (p = 0.037, η_p_^2^ = 0.141).

No differences were found between males and females weight changes and triglyceride and cholesterol levels. No levels were outside of the normal range.

### 2.7. Hepatic performance

GOT, GPT, GGT, ALP and albumin levels significantly decreased during ARI treatment (p = 0.001, η _p_ ^2^ = 0.249; p = 0.004, η _p_ ^2^ = 0.209; p = 0.001, η _p_ ^2^ = 0.224; p < 0.001, η _p_ ^2^ = 0.312; p < 0.001, η _p_ ^2^ = 0.307, respectively) (*Table* S3). Additionally, GGT levels were inversely proportional to DARI and ARI+DARI C_max_ (p = 0.050, η_p_^2^ = 0.116; p = 0.043, η_p_^2^ = 0.121). Likewise, ALP levels were dependent on ARI and ARI+DARI C_max_ (p = 0.042, η _p_ ^2^ = 0.121; p = 0.048, η _p_ ^2^ = 0.117).

OLA treatment produced a decline in GGT, bilirubin, ALP and albumin levels (p < 0.001, η_p_^2^ = 0.281; p = 0.045, η _p_ ^2^ = 0.123; p = 0.007, η _p_ ^2^ = 0.215; p = 0.004, η _p_ ^2^ = 0.285, respectively). All GOT, GPT, GGT, bilirubin and ALP levels normalized after discontinuing ARI or OLA treatment (*Table* S3).

No differences were found between males and females in GOT, GPT, GGT, ALP, bilirubin and albumin levels. Additionally, the changes in albumin levels differed between ARI and OLA treatment (p = 0.009, η_p_^2^ = 0.183).The changes in the rest of the parameters were not dependent on the treatment (ARI versus OLA). No levels were outside of the normal range.

### 2.8. Haematological parameters

Platelet count significantly decreased during ARI treatment (p < 0.001, η_p_^2^ = 0.361). Additionally, the prothrombin time increased and the prothrombin index decreased over time (p < 0.001, η_p_^2^ = 0.360; p < 0.001, η_p_^2^ = 0.410, respectively) (*Table* S4).

On the contrary, the leucocyte and platelet count decreased during OLA treatment (p = 0.004, η_p_^2^ = 0.217; p = 0.007, η_p_^2^ = 0.199, respectively). Similarly to ARI, the prothrombin index decreased over time (p = 0.006, η_p_^2^ = 0.237) (*Table* S4).

No differences were found between males and females in leucocyte, platelet, haemoglobin, red blood cell and haematocrit count and prothrombin time and index. No levels were outside of the normal range.

### 2.9. Thyroid function

Free T4 levels significantly increased after ARI treatment (p = 0.035, η _p_ ^2^ = 0.180). On the contrary, after OLA treatment, decreased levels were observed. However, this association did not reach the significant level (p = 0.230). Neither ARI, nor OLA had a significant effect on TSH levels (*Figure* 3). Nonetheless, a significant difference was found between ARI and OLA treatment in both free T4 and TSH levels (p = 0.010, η _p_ ^2^ = 0.267; p = 0.022, η _p_ ^2^ = 0.216). No differences were found between males and females in free T4 and TSH levels. No levels were outside of the normal range.

**Figure 3.**
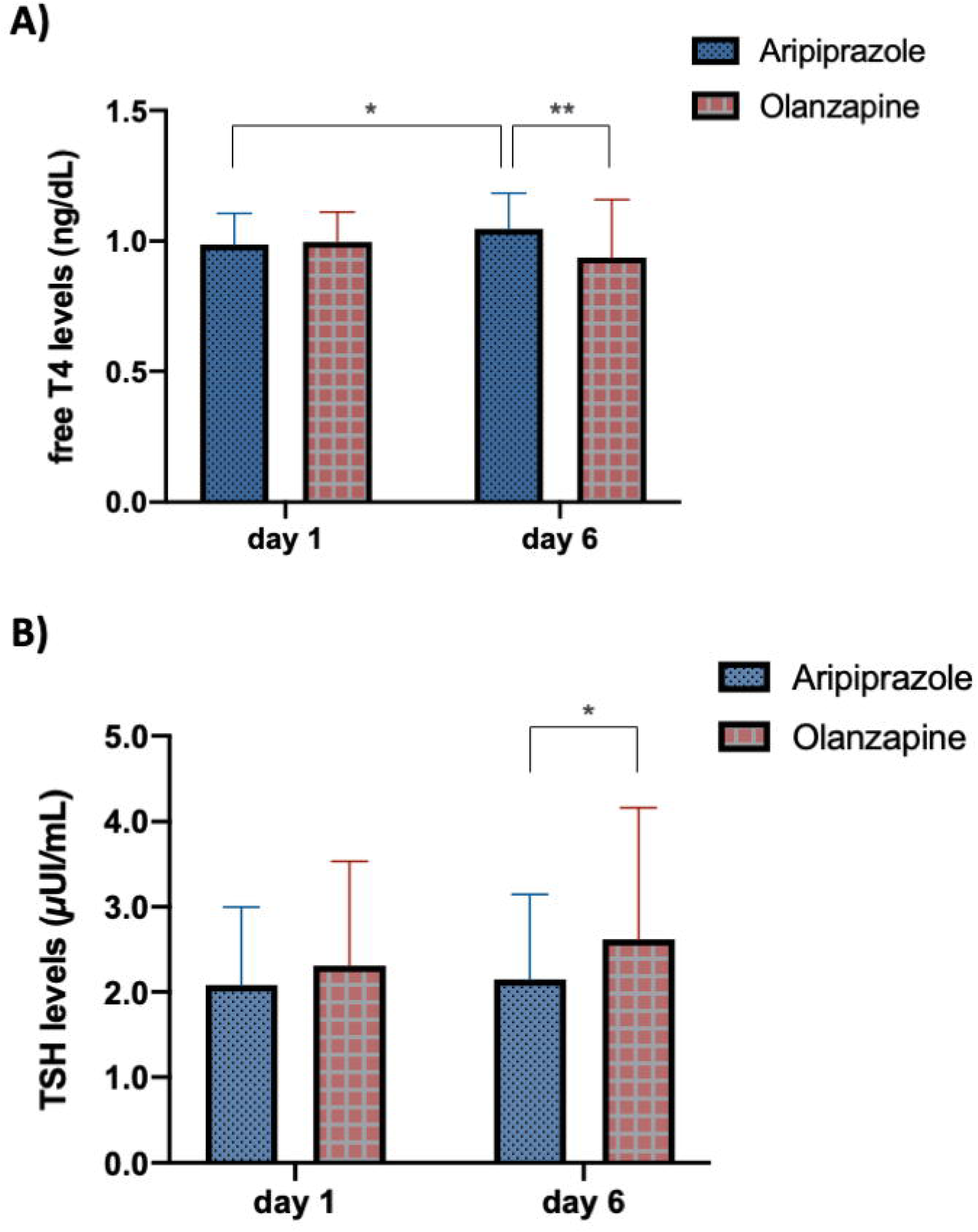
Free T4 and TSH concentrations after multiple dose administration of aripiprazole 10 mg and olanzapine 5 mg tablets. The values are shown as mean ± SD. *\* p*□< □0.05, *\*\* p*□< □0.01. ***Figure* 3A**. Free T4 concentrations after aripiprazole and olanzapine treatment. ***Figure* 3B**. TSH concentrations after aripiprazole and olanzapine treatment.

### 2.10. Renal function

Uric acid levels significantly decreased during ARI and OLA treatment (p < 0.001, η_p_^2^ = 0.324; p = 0.045, η_p_^2^ = 0.116) (*Table* S5). No differences were found between males and females in urea, creatinine and uric acid levels. Additionally, the changes in their levels were not dependent on the treatment (ARI versus OLA). The uric acid levels were not outside of the normal range.

The summary of the effects of ARI and OLA on all metabolic parameters is shown in *Table 3*.

**Table 3.** The effects of aripiprazole and olanzapine on all measured metabolic parameters.

|  | Aripiprazole | Olanzapine |
| --- | --- | --- |
| **Prolactin (ng/mL)\*** |  | ↑ |
| **C-peptide (ng/mL)** | ↑ | ↑ ns |
| **Insulin (mcU/mL)** | ↑ ns | ↑ ns |
| **C-peptide/insulin ratio** |  | ↑ |
| **HbA1c (%)** |  |  |
| **Glucose (ng/mL)** |  |  |
| **GTT (mg/dL)\*** | ↑ ns | ↑ |
| **Weight (kg)\*** | ↓ ns | ↑ ns |
| **Triglyceride (mg/dL)** | ↓ | ↓ |
| **Cholesterol (mg/dL)** | ↓ | ↓ |
| **GOT (U/L)** | ↓ |  |
| **GPT (U/L)** | ↓ |  |
| **GGT (U/L)** | ↓ | ↓ |
| **Total bilirubin (mg/dL)** |  | ↓ |
| **ALP (U/L)** | ↓ | ↓ |
| **Leucocytes (/μL (10<sup>3</sup>))** | ↓ ns | ↓ |
| **Platelets (/μL (10<sup>3</sup>))** | ↓ | ↓ |
| **Hemoglobin (mg/dL)** | ↓ ns | ↓ ns |
| **Red blood cells (/μL (10<sup>6</sup>))** | ↓ ns | ↓ ns |
| **Hematocrit (vol%)** | ↓ ns | ↓ ns |
| **Prothrombin time (sec)** | ↑ |  |
| **Prothrombin index (%)** | ↓ | ↓ |
| **Albumin (g/dL)\*** | ↓ | ↓ |
| **Free T4 (ng/dL)\*** | ↑ |  |
| **TSH (μUI/mL)\*** |  | ↑ ns |
| **Urea (mg/dL)** |  |  |
| **Creatinine (mg/dL)** |  |  |
| **Uric acid (mg/dL)** | ↓ | ↓ |

P ≤ 0.05 aripiprazole compared to olanzapine.
Abbreviations: HbA1c: haemoglobin A1c, GTT: glucose tolerance test, GOT: glutamate-oxaloacetate transaminase, GPT: glutamate-pyruvate transaminase, GGT: gamma-glutamyl transferase, ALP: alkaline phosphatase, free T4: free thyroxine, TSH: thyroid stimulating hormone, ns: non-significant.

## 3. Discussion

OLA caused prolactin elevation instantly after administering the first dose. Hyperprolactinemia is a common side effect of OLA treatment along with other atypical antipsychotics ^20^, what is produced by DRD2 blockage. Therefore, it causes loss of the dopaminergic prolactin inhibitory factor in the lactotroph cells in the anterior pituitary. Hence, antipsychotics with a greater D2 occupation index produce significant prolactin elevation ^21^. Our previous study revealed that prolactin levels significantly increase after administering a single dose of OLA ^22^. Our current study confirms that 5 multiple doses of OLA treatment also causes prolactin elevation.

Prolactin levels decreased after changing the therapy from other atypical antipsychotics - including OLA - to ARI ^23^. Our previous study showed that a single dose of ARI mildly increases prolactin levels compared to the controls ^13^. However, compared to OLA, no change was observed in prolactin levels after administration in the present study. Our study is the first to report a comparison between prolactin elevation induced by ARI and OLA in the same subjects. Therefore, the results can be considered reliable as the intraindividual variability is discarded. Schizophrenic patients usually receive several antipsychotic agents before ARI, therefore they are almost never drug-naïve ^23^. Our healthy subjects had not received any antipsychotic medication previously, hence no prior drug treatment could cause prolactin elevation. The current clinical practice recommends switching to ARI monotherapy in case of having high prolactin levels and if it does not appear to be normalized after 4 weeks of treatment, it should be discontinued ^24^.

Sex has a clear effect on prolactin concentrations, what was confirmed in several studies ^12,25^. Its levels tend to be higher in females than in males ^25^. Therefore, ARI and OLA effects on prolactin secretion were analysed both jointly and separately.

*HTR2C* rs17326429 and rs3813929, *COMT* rs4680, *DRD2* rs1800497, *DRD3* rs6280 and *ABCB1* rs1045642, rs1128503, rs2032582 and rs2235048 polymorphisms and CYP2D6 phenotypes were previously associated to prolactin levels after risperidone, quetiapine, clozapine, ARI or OLA treatment ^10,13,26,27^. We expected similar results as ARI is a partial agonist at DRD2 and at 5HT1A receptors and an antagonist at 5HT2A receptors while OLA is an antagonist at DRD2 and at 5HT2A and 2C receptors ^28,29^. In addition, ARI is metabolized by CYP2D6 and CYP3A4. However, ARI usually does not induce hyperprolactinemia ^6^, therefore a clear difference in prolactin levels would not be expected among phenotype groups. However, CYP3A PM subjects had significantly higher prolactin concentrations compared to the other phenotypes during ARI treatment. These subjects were under prolonged ARI exposure what could cause mild prolactin increase. Regarding *ABCB1*, subjects with rs10280101, rs12720067 and rs11983225 A-C-T haplotype had higher prolactin concentrations compared to those carrying the mutated alleles. This confirms the hypothesis that *ABCB1* polymorphisms and haplotypes might affect P-glycoprotein activity, therefore ARI brain availability and prolactin levels ^27^. Regarding OLA, *DRD3* rs6280 Ser/Ser subjects had lower prolactin levels compared to those carrying the Gly allele. Consequently, they may show higher DRD3 occupancy, thus dopamine can inhibit prolactin release ^6^. Previous findings imply that *DRD3* does not play a major role in OLA-induced prolactin secretion ^30^. Notwithstanding, we did not expect these results considering that none of the *DRD2* polymorphisms affected prolactin levels. More studies are needed to resolve the ambiguity.

C-peptide levels were significantly higher after ARI treatment. However, these levels were not significantly different between ARI and OLA. Clozapine, OLA, risperidone and sulpiride were associated with an increase in C-peptide levels in schizophrenic patients ^31^. In our study, OLA tended to increase its levels without reaching statistical significance; however, the C-peptide/ insulin ratio was higher after the 5 days treatment. This ratio is an indirect index of hepatic insulin clearance ^32^. The observed increase in the ratio may be due to the increase in hepatic insulin clearance therefore decreased insulin secretion ^33^. Teff *et al* with similar study design – 3 days of OLA treatment in healthy volunteers – found the contrary: a decrease was observed in the ratio what may imply an increase in insulin secretion ^34^. In the latter study, opposite to us, ARI did not cause elevation in C-peptide levels ^34^. High C-peptide levels can imply insulin resistance and finally can lead to type 2 diabetes, atherosclerosis and metabolic syndrome. Thus, it may serve as a biomarker to identify the risk to develop these diseases ^35^.

*COMT* rs4680 G/G and rs1330678 T carriers had higher increase in C-peptide levels after ARI treatment. *COMT* polymorphisms were previously associated with glycaemic function and type 2 diabetes, what can alter catecholamine production ^36^. *COMT* rs4680 A carriers achieved a significantly lower change in C-peptide levels compared to subjects with G/G genotype what is consistent with a previous study ^37^. Thus, A may be the protective allele against changes in glucose metabolism. ARI and OLA seem to have an effect on C-peptide levels, however, more studies are needed both in patients and healthy volunteers to confirm these findings.

OLA is associated with glucoregulatory abnormalities. The 5-HT1 antagonism may decrease the responsiveness of the pancreatic beta cells, thus reducing the secretion of insulin and causing hyperglycemia ^38^. In our study, basal glucose levels did not change during acute treatment, however, the GTT performed after treatment showed higher 1 h and 2 h glucose levels compared to the first day. These levels were significantly higher than during ARI treatment. Previous findings show the same association in patients undergoing chronic treatment and healthy volunteers with acute treatment ^39,40^. *UGT1A1* rs887829 C/C homozygotes had higher basal glucose levels and also higher glucose levels in GTT after 1 h and 2 h on day 6 compared to the first day. OLA is metabolized predominantly by the UGT enzyme family, but clear evidence was found only for UGT1A4 ^41^. Based on our results, T allele carriers may be under prolonged OLA exposure and therefore show higher glucose concentrations. This result is confirmed by our previous study as this polymorphism affected OLA pharmacokinetics ^42^.

ARI and OLA tended to increase insulin levels. In addition, *BDNF* rs6265 C/C subjects showed higher insulin levels compared to the other genotypes after ARI administration and in CYP3A PMs compared to the other phenotypes after OLA administration. In a previous study the *BDNF* rs6265 polymorphism did not affect insulin levels during chronic risperidone and OLA treatment ^43^. Based on our knowledge, our study is the first to report this relationship with ARI. C/C subjects may have more predisposition to develop high insulin levels and finally insulin resistance during ARI treatment.

It is not completely understood how antipsychotics cause weight gain, but 5-HT2C and 5-HT1A receptors, histamine H1 receptor and DRD2 presumably play a role ^44^. However, OLA pharmacology is not the only factor to affect weight gain; the diet and activity level may also play a role. Weight increases rapidly within the first 6 weeks of OLA treatment and patients continue to gain weight ^45^. Based on our knowledge, our study is the first to report OLA-related weight gain during only 5 days of treatment. ARI did not induce weight gain in the same volunteers, what strengthens our results as we can discard the effect of the diet. *HTR2C* polymorphisms are clearly linked to susceptibility to gain weight with antipsychotics ^46^. The *HTR2C* rs1414334 polymorphism was widely analysed and the C allele was associated to OLA, clozapine and risperidone-induced weight gain and metabolic syndrome ^47^, what we confirm in our study.

Based on current knowledge, OLA, but not ARI, increments triglyceride and cholesterol levels in chronic treatment ^48^. Moreover, in a previous study, after administering 3 daily doses of OLA to healthy volunteers, the cholesterol and the triglyceride levels were higher ^40^. These results suggest that OLA may have acute adverse effects on lipid profiles as well. However, in our study, both triglyceride and cholesterol levels decreased during ARI and OLA treatment. The observed decrease could be due to the low carbohydrate diet during their stay ^49^. It could explain why triglyceride and cholesterol levels recovered by the safety visit (10 days after discontinuing the treatment).

CYP3A PMs showed a greater decrease in triglyceride levels during ARI treatment compared to the other phenotypes. PMs could have a prolonged ARI exposure, therefore higher effect on the triglyceride levels as ARI is metabolized by CYP3A ^5^. Furthermore, *APOC3* rs4520 C/C homozygotes had higher triglyceride concentrations after OLA administration than T allele carriers, what is consistent with a previous study ^50^. Polymorphisms in this gene influence serum or plasma triglyceride levels as the APOC3 protein raises plasma triglyceride levels by the inhibition of lipoprotein lipase, stimulates low-density lipoprotein secretion and intestinal triglyceride trafficking modulation ^51^. The *HTR2A* rs6314 C/C homozygotes had greater cholesterol levels during ARI therapy. In a previous study the contrary was found: T carriers had higher cholesterol levels. However, Korean population when compared to Caucasians, have a lower frequency of *HTR2A* C allele (approximately 0.515), therefore the frequency variation among different ethnic groups could cause this variation ^52^. Our study is the first to report differences between *HTR2A* rs6314 alleles in cholesterol level changes during antipsychotic therapy.

In a previous study, ARI elevated mildly, while OLA elevated greatly the transaminase levels (eg. GOT and GPT) ^53^. ARI effects on GGT and ALP levels were not reported to date. However, OLA was reported to increase GGT, ALP and bilirubin levels ^53^. Based on our knowledge, our study is the first to report changes in hepatic enzyme and bilirubin levels during short time antipsychotic treatment in healthy volunteers. In our study, GOT, GPT, GGT and ALP levels significantly decreased during ARI treatment while GGT, bilirubin and ALP levels significantly decreased during OLA treatment. The observed decrease could be explained by the low carbohydrate diet during their stay ^49^. Additionally, the albumin levels also decreased during ARI and OLA treatment alike risperidone and clozapine, what suggest that these drugs have a negative impact on serum antioxidant protection ^54^. In addition, none of these levels were outside of the reference range.

OLA may cause leukopenia ^55^, thrombocytopenia ^56^ and thromboembolism ^57^. ARI only causes these conditions when co-administered with other CYP2D6 substrates ^58^. Our study confirms that both antipsychotics cause significant decrease in platelet count and OLA additionally induces a decrease in leucocyte count. The current study is the first to report that these alterations start immediately after starting treatment, although none of these levels were outside of the reference range.

The free T4 levels significantly increased after ARI treatment. ARI drug label states that it can induce both hypo,- and hyperthyroidism. Nevertheless, the underlying mechanism is currently unknown ^4^. In studies with quetiapine, only free T4 changes were detected, and not TSH ^59^, similar to our results. Additionally, compared to the ARI group, the OLA group had higher TSH and lower free T4 levels after treatment. OLA was associated with lower free T4 and higher TSH levels in patients compared to healthy controls in a previous study ^60^. Our study is the first to report increase in free T4 levels after ARI treatment in healthy volunteers.

Atypical antipsychotics can increase the risk to develop chronic kidney diseases through the elevation of urea and creatinine levels ^61,62^. After the acute treatment with ARI and OLA we could not see this effect. Uric acid levels decreased during haloperidol ^63^, but not risperidone or clozapine treatment ^54^. In the current study, we observed that both ARI and OLA reduced its levels during treatment, but the levels were normalized after discontinuing the drugs. Uric acid is one of the principal antioxidants in the human plasma, therefore its low levels may cause oxidative stress ^54^. Based on the authors’ knowledge, this is the first study to analyse uric acid alterations in acute ARI and OLA treatment.

## 4. Study limitations

The most significant limitation of our study is the low sample size. Therefore, it is of importance to interpret these results with caution. Our study should be repeated in healthy volunteers to increase the sample size as well as in schizophrenic patients to demonstrate the clinical utility of these results. Despite of applying the Bonferroni post hoc test to each analysis, some of our results could be false positives due to the high number of analysed variables. Moreover, neither ARI, nor OLA reached steady state during 5 days of treatment. Both could have had a greater effect on metabolism if they had reached steady state. Nevertheless, we had very well controlled conditions what can reduce the influence of other factors.

## 5. Conclusions

OLA caused significant prolactin elevation, but not ARI. CYP3A phenotype and *ABCB1* and *DRD3* genes affected the prolactin levels. ARI caused C-peptide elevation and were dependent on *COMT* genotypes while the C-peptide/ insulin ratio was higher after OLA treatment. Glucose levels in GTT were higher after 5 doses of OLA and were influenced by *UGT1A1* genotypes. Insulin levels did not change but were dependent on *BDNF* genotypes and CYP3A phenotypes. OLA caused weight gain, what was influenced by *HTR2C* alleles. Triglyceride and cholesterol levels decreased during ARI and OLA treatment. Triglyceride levels were variable based on CYP3A phenotypes and *APOC3* genotypes. Cholesterol levels were dependent on *HTR2A* genotypes. ARI induced an increase in free T4 levels. Both ARI and OLA reduced uric acid levels. Both antipsychotics have significant metabolic effects in acute treatment. However, we can confirm that ARI has a more benign metabolic profile.

## 6. Methods

### 6.1. Study Population and Design

The study population comprised 24 healthy volunteers (12 men and 12 women) who were enrolled in a phase I, 5 daily oral doses, open-label, randomized, crossover, two-period, two-sequence, single-centre, comparative clinical trial. This sample size was considered sufficient to detect a clinically important difference in metabolism. Ten mg/day ARI tablets or 5 mg/day film-coated OLA tablets were administered (Laboratorios Alter, Madrid, Spain). Block randomization was used to assign the treatment to each volunteer on the first day ^64^. The whole duration of the hospitalization was 5 days since 1 h before the first dose until 24 h after the administration of the last dose. The drug was administered at 09:00 h each day under fasting conditions. After a washout period of 28 days, each volunteer received the opposite drug that they had received in the first period. The random allocation sequence, the recruitment of participants and their assignment to interventions were performed by investigators of the Clinical Trials Unit.

The clinical trial was performed at the Clinical Trials Unit of our hospital. The protocol was approved by its Research Ethics Committee authorized by the Spanish Drugs Agency and the study was performed under the guidelines of Good Clinical Practice and the Declaration of Helsinki (Clinical Trial registry name, registration number, URL and date: TREATMENT-HV, EUDRA-CT: 2018-000744-26, https://eudract.ema.europa.eu/, April 15, 2018). All subjects signed the informed consent before inclusion and were free to withdraw from the study at any time.

The inclusion criteria were the following: male and female volunteers between 18 and 65 years old; free from any known organic or psychiatric conditions; normal vital signs and electrocardiogram (ECG); normal medical records and physical examination; no clinically significant abnormalities in haematology, biochemistry, serology and urine tests.

The exclusion criteria were the following: received prescribed pharmacological treatment in the last 15 days or any kind of medication in the last 48 hours prior to receiving the study medication; body mass index (BMI) outside the 18.5-30.0 kg/m^2^ range; history of drug allergy; having galactose intolerance, Lapp lactase deficiency or glucose-galactose malabsorption; suspected consumers of controlled substances; smokers; daily alcohol consumers and/or experienced acute alcohol poisoning the previous week; donated blood last month; pregnant or breastfeeding women; investigational drug study participants in the previous 3 months and subjects unable to follow instructions or collaborate during the study.

### 6.2. Pharmacokinetic Analysis

Twenty-two blood samples were collected until 10 days after the last dose in EDTA K2 tubes for pharmacokinetic assessments during each period of the study. Samples were centrifuged at 3500 rpm (1900 G) for 10 minutes and then the plasma was collected and stored at -80 °C until the determination of plasma concentrations. Plasma concentrations of ARI, DARI and OLA were quantified by a high-performance liquid chromatography tandem mass spectrometry (HPLC-MS/MS) method developed and validated in our laboratory ^65^.

The pharmacokinetic parameters were calculated by noncompartmental analysis by Phoenix^®^ WinNonlin^®^ (version 8, Pharsight, Mountain View, CA, USA). The peak concentration (C_max_) was obtained directly from the original data. The area under the plasma concentration-time curve from time zero to the last observed time point (AUC_last_) was calculated using the trapezoidal rule. The AUC from time zero to infinity (AUC_inf_) was determined as the sum of the AUC_last_ and the extrapolated area beyond the last plasma concentration. The terminal rate constant (ke) used for the extrapolation was determined by regression analysis of the log-linear part of the concentration-time curve. The AUC and C_max_ were adjusted for dose and weight (AUC/dW and C_max_/dW, respectively) and were logarithmically transformed for statistical analysis ^18^.

### 6.3. Biochemical and Haematological Analyses

The biochemical and haematological analyses were carried out by Eurofins Megalab S.A. (Madrid, Spain). All subjects underwent the oral glucose tolerance test (GTT) on days 1 and 6 with 75 LJg of oral anhydrous glucose dissolved in 250 LJmL water. Glucose, triglyceride, bilirubin, glutamate-oxaloacetate transaminase (GOT), glutamate-pyruvate transaminase (GPT), gamma-glutamyl transferase (GGT), albumin, alkaline phosphatase (ALP), uric acid, urea and creatinine concentrations were measured spectrophotometrically and the samples were collected at screening and on days 3, 6 and 15. Cholesterol levels were analysed by enzymatic colorimetric method at screening and days 3, 6 and 15. Prolactin levels were analysed on days 1, 3 and 5 before and after dosing and days 6 and 15 and C-peptide, insulin, thyroid stimulating hormone (TSH) and free thyroxine (T4) concentrations were quantified on days 1 and 6 by Enzyme-Linked ImmunoSorbent Assay (ELISA). Hemoglobin A1c (HbA1c) was measured on days 1 and 6. Haematocrit, platelet, leucocyte, haemoglobin and red blood cell (RBC) counts were measured by flow cytometry at screening and on days 3, 6 and 15. Finally, prothrombin time was determined by coagulometry at screening and on days 3, 6 and 15.

### 6.4. Genotyping

DNA was extracted from 1 mL of peripheral blood samples using a MagNA Pure LC DNA Isolation Kit in an automatic DNA extractor (MagNa Pure® System, Roche Applied Science, Indianapolis, Indiana). Thenceforth, it was quantified spectrophotometrically in a NanoDrop® ND-1000 Spectrophotometer (Nanodrop Technologies, Wilmington, Delaware, USA) and the purity of the samples was measured by the A_260/280_ absorbance ratio.

The samples were genotyped with TaqMan assays using the OpenArray platform on a QuantStudio 12K Flex instrument (Thermo Fisher Scientific, Waltham, Massachusetts, USA). The genotyping array included 120 SNPs, whereof the following 51 were analysed in 18 genes due to their importance in the metabolism and the mechanism of action of ARI and OLA: *CYP1A2* *1C (rs2069514), *1F (rs762551), *1B 5347T>C (rs2470890), *CYP2D6* *3 (rs35742686), *4 (rs3892097), *6 (rs5030655), *7 (rs5030867), *8 (rs5030865), *9 (rs5030656), *10 (rs1065852), *14 (rs5030865), *17 (rs28371706), *41 (rs28371725), *CYP3A4* *22 (rs35599367), rs55785340, rs4646438, *CYP3A5* *3 (rs776746), *6 (rs10264272), *ABCB1* C3435T (rs1045642), G2677 T/A (rs2032582), C1236T (rs1128503), rs3842, 1000-44G>T (rs10276036), 2895+3559C>T (rs7787082), 330-3208C>T (rs4728709), 2481+788T>C (rs10248420), 2686-3393T>G (rs10280101), 2320-695G>A (rs12720067), 2482-707A>G (rs11983225), 2212-372A>G (rs4148737), *ADRA2A* rs1800544, *APOC3* rs5128, rs4520, *APOA5* rs662799, *BDNF* Val66Met (rs6265), *COMT* rs4680, rs13306278, *DRD2* TaqIA (rs1800497), 957C>T (rs6277), -141 Ins/Del (rs1799732), *DRD3* Ser9Gly (rs6280), *HTR2A* T102C (rs6313), C1354T (rs6314), rs7997012, *HTR2C* - 759C/T (rs3813929), -697G/C (rs518147), rs1414334, *LEP* rs7799039, *LEPR* rs1137101, *OPRM1* rs1799971 and *UGT1A1* rs887829 ^18^.

Results were analysed within both the QuantStudio(tm) 12K Flex and Thermo Fisher Cloud softwares (Thermo Fisher Scientific, Waltham, Massachusetts, USA). Finally, a matrix of genotypic calls was exported for each polymorphism.

The copy number variations (CNVs) in the *CYP2D6* gene were determined with the TaqMan^®^ Copy Number Assay (Assay ID: Hs00010001_cn; Thermo Fisher Scientific, Waltham, Massachusetts, USA) which detects a specific sequence on exon 9. The samples were run in the same instrument.

Whereas the *CYP2D6* *29 (rs16947) polymorphism was not included in the array, it was genotyped with the same instrument using individual TaqMan^®^ probes. Additionally, the *CYP3A4* *20 (rs67666821) polymorphism was genotyped by the KASPar SNP Genotyping System (LGC Genomics, Herts, UK). The ABI PRISM 7900HT Sequence Detection System (Thermo Fisher Scientific, Waltham, Massachusetts, USA) was used for fluorescence detection and allele assignment ^66^.

### 6.5. Statistical analysis

The statistical analysis was performed with the SPSS 24.0 software (SPSS Inc., Chicago, Illinois, United States). P values lower than 0.05 were considered statistically significant. The Hardy-Weinberg equilibrium was estimated for all genetic variants. Deviations from the equilibrium were detected by comparing the observed and expected frequencies using a Fisher exact test based on the De Finetti program (available at http://ihg.gsf.de/cgi-bin/hw/hwa1.pl). The metabolic parameters were analysed by repeated measures ANOVA. Pharmacokinetic parameters and polymorphisms were analysed as covariates. MANOVA was used to study factors related to all metabolic data and pharmacokinetic variables. The Bonferroni post-hoc test for multiple comparisons was applied for each analysis.

*CYP2D6* *3, *4, *5, *6, *7, *8, *9, *10, *14, *17, *29 and *41 were classified in phenotypes based on the functionality of the alleles ^67,68^. *CYP3A4* *2, *20, *22, and *CYP3A5* *3 and *6 genotypes were merged into a CYP3A phenotype based on Sanchez Spitman et al ^69^. *CYP1A2* *1C, *1F and *1B variants were also merged into a phenotype as previously reported ^70^. *ABCB1* variants were merged into haplotypes: 0-8 mutated alleles were assigned to group 1, 9-12 mutated alleles were assigned to group 2 and 13-17 mutated alleles were assigned to group 3. Another *ABCB1* haplotype was assembled by only considering C3435T, G2677T/A and C1236T polymorphisms due to greater impact on the transporter’s activity or expression levels ^71^. Zero or one mutated allele carriers were assigned to group 1, carriers of 2 or 3 mutated alleles were assigned to group 2 and bearing 4, 5 or 6 mutated alleles were assigned to group 3. *COMT* rs13306278 and rs4680 polymorphisms were merged into a haplotype: carrying no mutant allele was assigned as wild-type, carrying one mutant allele was considered as heterozygous while bearing more than one mutant allele was considered as mutant.

## Data availability

All data generated or analysed during this study are included in this published article (and its Supplementary Information files). Clinical Trial registry name, URL and registration number: TREATMENT-HV, EUDRA-CT: 2018-000744-26, https://eudract.ema.europa.eu/.

## Acknowledgements

The authors are grateful to the volunteers as well as the effort of the staff of the Clinical Trials Unit of La Princesa University Hospital, especially to Samuel Martín and Alejandro de Miguel-Cáceres for sample processing and to Daniel Romero-Palacián and María J. Hernández for safety evaluations.

## Author Contributions

Wrote Manuscript: DK, MSR, PZ, SA, FAS; Designed Research: DK, FAS, MSR, GM, MR, DO; Performed Research: DK, MR, GM, FAS, MN; Analysed Data: DK, SA, ESM, EPS.

